# Exceptional fossil preservation and evolution of the ray-finned fish brain

**DOI:** 10.1101/2022.06.04.492470

**Authors:** Rodrigo T. Figueroa, Danielle Goodvin, Matthew A. Kolmann, Michael I. Coates, Abigail M. Caron, Matt Friedman, Sam Giles

## Abstract

Brain anatomy provides key evidence for ray-finned fish relationships^1^, but two key limitations obscure our understanding of neuroanatomical evolution in this major vertebrate group. First, the deepest branching living lineages are separated from the group’s common ancestor by hundreds of millions of years, with indications that aspects of their brain morphology–like other aspects of their anatomy^2,3^–are specialised relative to primitive conditions. Second, there are no direct constraints on brain morphology in the earliest ray-finned fishes beyond the coarse picture provided by cranial endocasts: natural or virtual infillings of void spaces within the skull^4–8^. Here we report brain and cranial nerve soft-tissue preservation in †*Coccocephalichthys wildi*, a ∼319-million-year-old (Myr) ray-finned fish. This oldest example of a well-preserved vertebrate brain provides a unique window into neural anatomy deep within ray-finned fish phylogeny. †*Coccocephalichthys* indicates a more complicated pattern of brain evolution than suggested by living species alone, highlighting cladistian apomorphies^9^ and providing temporal constraints on the origin of traits uniting all extant ray-finned fishes^9–11^. Our findings, along with a growing set of studies in other animal groups^12–16^, point to the significance of ancient soft tissue preservation in understanding the deep evolutionary assembly of major anatomical systems outside of the narrow subset of skeletal tissues^17–20^.

## MAIN

Actinopterygian (ray-finned fish) brains display anatomical innovations not seen in other vertebrates, most notably a forebrain that grows in a unique way: through eversion of the dorsal walls of the telencephalon, rather than evagination of its lateral walls^21,22^. This results in a forebrain formed of two solid hemispheres that do not enclose a ventricle^23^. Brain anatomy therefore provides important evidence for the monophyly and interrelationships of ray-finned fishes, a major radiation containing roughly half of all vertebrate species^24^. Brain anatomy in non-teleost fishes is limited to a handful of examples, reflecting the low diversity of the deepest extant branches of the ray-finned fish tree of life. Fossils provide limited constraints on brain structure deep in actinopterygian phylogeny. In contrast to teleosts, where the inner walls of the neurocranium are generally widely separated from most parts of the brain, the contours of the cranial endocavity (endocast) in non-teleosts appear to capture some aspects of neural anatomy^25^. For over a century, rare natural endocasts^4,26^ and a handful of models from physical tomography^5,27,28^ provided the only constraints on brain structure in early ray-finned fishes. The recent widespread application of computed tomography yields a greater number of examples spanning the very deepest branches of the actinopterygian tree^7^ to the teleost and holostean stems^25,29^, and several groups in between^6,30,31^. These provide information on gross morphological patterns of actinopterygian brain evolution and represent an important source of characters for phylogenetic analysis^6,26^. However, there are significant disconnects between our understanding of neural anatomy in fossil species, where information derives exclusively from the endocavity, and living forms, where only brain anatomy is well documented. This stems from two practical limitations: the low preservation potential of brain tissues in the fossil record combined with a poor understanding of endocavity anatomy in living taxa. Consequently, key evolutionary steps that preceded the origin of living actinopterygian brains remain unknown.

Although rare, there is a growing record of preserved neural tissue in fossils. Palaeozoic arthropods provide the most examples^12–16^, although there is a solitary report of a preserved brain in a Carboniferous cartilaginous fish allied to extant ratfishes^19^. Here we report an exceptionally preserved brain and associated cranial nerves in the Pennylvanian (Bashkirian; ∼319 Myr) ray-finned fish †*Coccocephalichthys wildi*, representing the first known fossil example for actinopterygians. Current analyses place this taxon outside the group containing all living species^29^. Details of brain structure in †*Coccocephalichthys* therefore bear on interpretations of neural morphology during the early stages of evolution in a principal lineage of backboned animals. Using μCT of fossil material in concert with diceCT imaging of extant species^32^, we provide a revised picture of brain evolution in bony fishes.

### Description

#### Endocast and otoliths

The endocast of †*Coccocephalichthys*, like that of other Palaeozoic actinopterygians, is clearly differentiated into areas that appear to correspond to regions of the brain (Fig. 1a). It agrees most closely with that described for *Lawrenciella*^28,33^. Only a single pair of otoliths, filling the saccular chamber, are preserved (Fig. 1b,d). These are large and teardrop shaped in lateral view, similar to those reported in some other Palaeozoic and early Mesozoic actinopterygians^34^. Their mesial and lateral surfaces are slightly convex and concave, respectively.

**Fig. 1:**
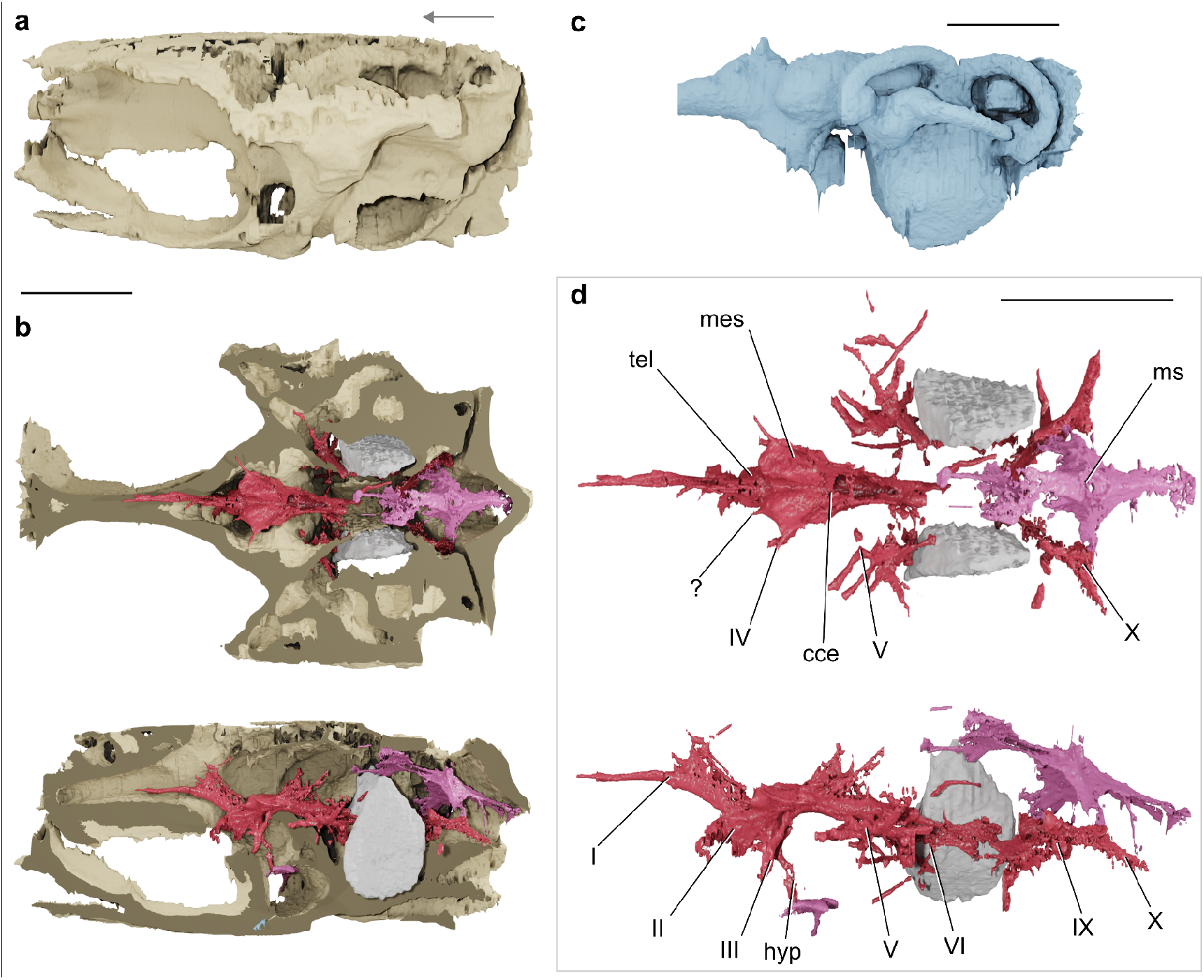
Neurocranium, endocast, otoliths and preserved brain of †*Coccocephalichthys wildi* (MM W.12451). **a**, Neurocranium in left lateral view. **b**, cutaways of neurocranium in dorsal (top) and left lateral (bottom) views showing brain and otoliths *in situ*. **c**, endocast in left lateral view. **d**, the brain and associated preserved soft tissues in dorsal (top) and lateral (bottom) views, with left otolith removed in the latter for clarity. cce, corpus cerebelli; hyp, hypophysis; mes, mesencephalon; ms, myelencephalic sheet; tel, telencephalon; I, olfactory nerve; II, optic nerve; III, oculomotor nerve; V, trigeminal nerve; VI, abducens nerve; IX, glossopharyngeal nerve; X, vagus nerve. Scale bars = 5 mm. Arrow indicates anterior for all panels.

#### Overall preservation of the brain

Within the cranial cavity lies a symmetrical object that is denser than the surrounding matrix (Fig. 2, Extended Data Figs. 1–5). It extends from the level of the orbit to the oticooccipital fissure. It comprises three principal structures: a central, hollow body that lies on the midline; ramifications on either side of the central body that are in some cases are clearly associated with endoskeletal nerve foramina; and a diamond-shaped sheet that lies posterodorsal to the other elements. The central body includes three regions: a long, narrow anterior extension; a swollen middle region comprising a horizontal plate with two dorsal hemispheres and a ventral outgrowth; and flattened posterior tube with a slit-like opening on the dorsal midline. Based on preservational style^19^ and comparison with neural features in extant jawed vertebrates (Fig. 2, Extended Data Fig. 3), we interpret this structure as a preserved brain. The three regions described above roughly correspond to the forebrain, midbrain and hindbrain, respectively, and collectively occupy around 5% of the endocranial volume.

**Fig. 2:**
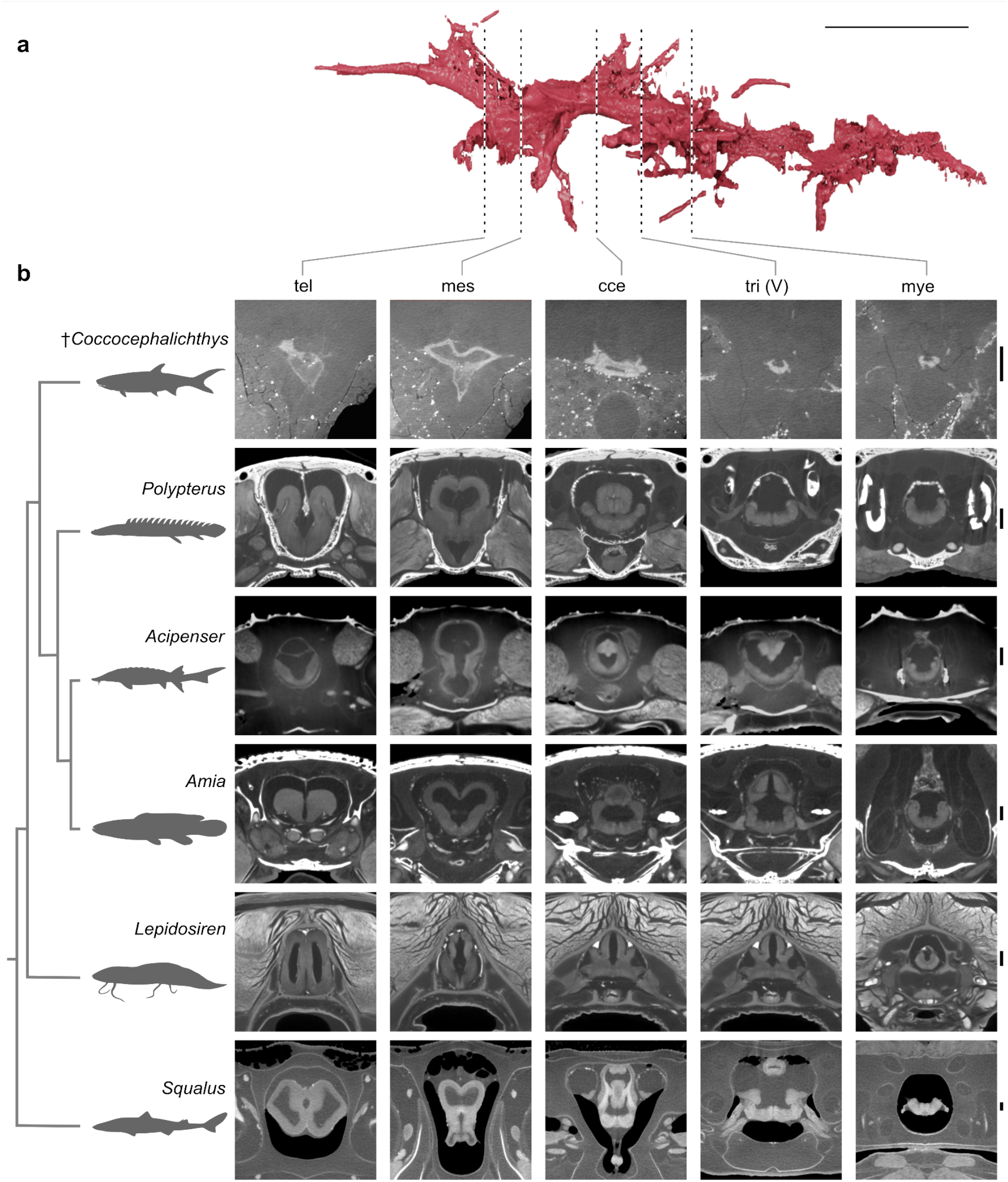
Anatomical correspondence between preserved brain of †*Coccocephalichthys wildi* and those of extant fishes. **a**, three-dimensional rendering of the brain of †*Coccocephalichthys* in left lateral view. Scale bar = 10 mm. **b**, transverse sections through the brains of †*Coccocephalichthys* and selected jawed fishes from diceCT data. cce, corpus cerebelli; mes, mesencephalon; mye, myelencephalon; tel, telencephalon; tri (V), trigeminal nerve; ?, unidentified midbrain feature. Silhouettes of extant taxa from phylopic2 (*Squalus*, Ignacio Contreras; *Lepidosiren*, Roberto Diaz Sibaja; *Acipenser*, Maija Karala, *Amia*, no copyright; *Polypterus*, no copyright). †*Coccocephalichthys* silhouette original based on ‘*Palaeoniscus’*.^59^ Scale bar = 1 mm.

#### Forebrain

The forebrain, comprising the olfactory bulbs, telencephalon and diencephalon, lies anterior to, and is considerably smaller than, the midbrain (Fig. 1). An elongate, slender extension anterior to the telencephalic body represents the olfactory nerve, but the olfactory bulbs are difficult to identify. The olfactory nerve extends to the midpoint of the orbit as a single tract before dividing anteriorly. A dorsal sheet extends into the pineal chamber posterior to the divergence of the olfactory tract from the telencephalon. This structure may represent the remnants of the velum transversum. Thin filaments connect the anterior and posterior margins of this sheet to the inner walls of the endocranial chamber, and paired anterior cerebral veins exit from its base. The body of the telencephalon is formed by two small, paired swellings separated by a median septum that is visible most clearly anteriorly and posteriorly (Extended Data Fig. 5). The swellings are moderately expanded laterally, giving the telencephalon an ellipsoidal profile in axial section (Fig. 2; Extended Data Fig. 5). Each swelling is hollow and encloses a large ventricular space, indicating that the forebrain is evaginated as in sarcopterygians and chondrichthyans^35–37,21^. By contrast, all living ray-finned fishes possess an everted telencephalon^9,21,38–40^ (Fig. 2B). We interpret an additional tissue layer dorsal to the telencephalon as part of the meningeal tissue of the forebrain.

There is no clear boundary dividing the telencephalon and diencephalon. A moderate expansion posteroventral to the telencephalon corresponds with an ellipsoidal ventricle within the main body of the brain, indicating the presence of partially developed hypothalamic inferior lobes (Extended Data Fig. 4,5). The lobes are visible in cross-section as small ellipsoid structures of a slightly denser material than the matrix, but less dense than the external brain wall. The right lobe is apparent externally on the right side of the brain as a low swelling. A slender and ventrally elongated hypophysis extends from behind the hypothalamus. It leads to a differentiated distal portion in contact with the buccohypophysial canal, and a posterior expansion associated with the saccus vasculosus. The ventricular space within each hypothalamic inferior lobe is connected with that of the hypophysis (the diencephalic ventricle) via a narrow canal, named the lateral hypothalamic recess^41^. The morphology of this structure in †*Coccocephalichthys* is similar to that of *Amia* (Extended Data Fig. 6).

#### Midbrain

The mesencephalic lobes, the dorsal surfaces of which comprise the optic tectum, are well-developed and oval in dorsal view (Fig. 1). The lobes are connected posteriorly, level with the cerebellar region, and diverge anteriorly. Two nerves emerge from the surface of the mesencephalon: a narrow, anterodorsally directed trochlear (IV) nerve; and a stout, anteroventrally directed oculomotor (III) nerve, which dichotomises within the braincase wall and enters the orbital cavity through two foramina. A third feature, which leaves the anterior margin of the midbrain, is of unclear identity. The optic chiasma is preserved on the anteroventral surface of the mesencephalon, along with the proximal portions of the optic (II) nerves. These extend and diverge beyond the external margin of the midline optic foramen.

Ventricles are apparent in sections through the midbrain (Fig. 2, Extended Data Figs. 1,2). The second (mesencephalic) ventricle mirrors the shape of the optic tectum, and is V-shaped in axial section and U-shaped in horizontal section. There does not appear to be either a torus longitudinalis or torus semicircularis within the second ventricle. Anteriorly, the mesencephalic ventricles connect to a tube-like ventricle that opens at the roof of the diencephalic region of the brain. Posteriorly the mesencephalic ventricles contact the fourth ventricle through a narrow tube-shaped connection.

#### Hindbrain

Few features of the hindbrain are preserved. The anteriormost portion of the hindbrain is developed as small rounded cerebellar auricular lobes, which are separated by the posterior limits of the mesencephalic lobes (Fig. 1). Posterior to these lies the recessus lateralis of the fourth ventricle, which is continuous with a thin, dorsally-extensive rhombencephalic tela choroidea. The cerebellar corpus is barely developed. The fourth ventricle is open dorsally. It is anteroposteriorly elongate and circular in axial section, and lies ventral to the mesencephalic ventricle (Fig. 2, Extended Data Figs. 1,2). A cerebral aqueduct connecting the second and fourth ventricles is not apparent. The internal walls of the fourth ventricle lack pronounced ridges, but it is difficult to say whether this is original or a taphonomic artifact. Two thin, posteroventrally directed branches of the abducens (VI) nerve leave the ventral surface of the brain level with posterior margin of the fourth ventricle. More ventrally, an additional branch extends from the saccular chamber in the direction of the posterior myodome. Due to the position and path of this branch, we identify it as a distally diverging branch of the abducens nerve.

The trigeminofacial nucleus and associated nerves are separated from the body of the hindbrain, and this is presumably a taphonomic artifact (Fig. 1). The trigeminofacial complex on the right of the specimen appears to be associated with the alar wall of the rhombencephalon, which has pulled away from the remainder of the hindbrain. Nerve branches located at the front of this complex are enclosed within skeletal canals and can thus be identified most readily by comparison with endocasts described for Palaeozoic actinopterygians, although we caution that this nomenclature needs review in comparison to nerve patterns in extant non-teleost actinopterygians. Two stout nerves emerge anterolaterally from the front of this complex, the most anterior of which enters the canal identified as that for the trigeminal (V) nerve, and the more posterior one the lateralis branch of the facial (VIIlat) nerve. A third nerve, which leaves the complex anteroventrally, enters the canal for the main branch of the facial (VII) nerve. More posteriorly, a series of nerves are associated with the inner ear and otolith, and most likely correspond to branches of the octavolateralis (VIII) nerve. The anterior branch of the anterior ramus of the octavolateralis extends some way dorsally into the anterior ampulla, with the posterior branch of the anterior ramus entering the utriculus. A posteroventral branch contacts the anterior margin of the otolith within the saccular cavity. Two to three additional rami attach to the medial margin of the otolith, and further branches may be present posteriorly.

A diamond-shaped sheet lies posterodorsal to the preserved portion of the brain, in close association with the roof of the endocranial cavity (Fig. 1, Extended Data Fig. 3). This structure is in a similar position to the meninx primitiva, modified to a cisterna spinobulbularis in *Polypterus*^42,43^, and a myelencephalic gland in other early ray-finned fishes^44^. The dorsal surface of the tissue sheet bears a medially located opening surrounded by a thin layer of tissue that extends as a tube toward the posterodorsal fontanelle of the neurocranium. The vagus (X) nerve lies ventral to this sheet, extending posterolaterally to exit from the braincase via the oticooccipital fissure. Anterior to the vagus nerve root, the glossopharyngeal nerve extends laterally towards the endocranial wall.

## Discussion and Conclusions

### Correspondence between brains and endocasts

It has been widely assumed that there is close correspondence between brain and endocast shape in early ray-finned fishes^4,6,26,42^. However, the brain as preserved in †*Coccocephalichthys* does not closely conform to the inner surface of the endocavity (Fig. 1, Extended Data Fig. 1). It seems likely that the brain has contracted to some degree during preservation, but the fact that many cranial nerves both connect with the brain itself and extend out of their neurocranial foramina places a limit on the degree of shrinkage.

There is also a clear positional match between regions of the preserved brain and areas of the endocavity hypothesised to accommodate them. Living ray-finned fishes show varying degrees of correspondence between brain and endocast morphology^46,47^ (Fig. 3), although in no case does the brain completely fill the endocavity in a way comparable to lungfishes and some tetrapods^48– 51^. This does not invalidate endocasts as sources of characters or information about neuroanatomy, but stresses that the anatomy of brains and endocavities should not be treated as interchangeable.

**Fig. 3:**
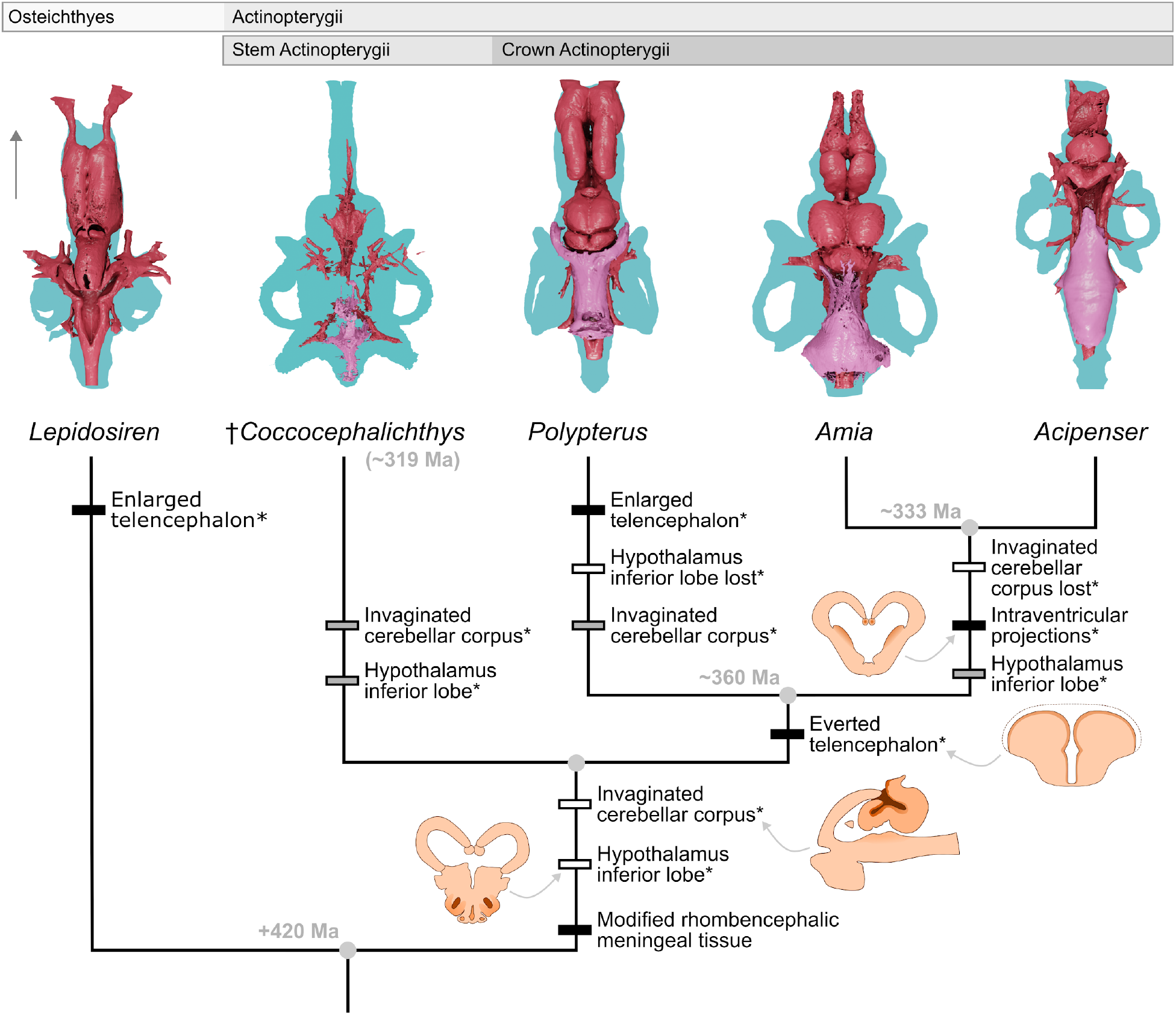
Major anatomical transformations in actinopterygian brain structure illuminated by †*Coccocephalichthys*. Branch labels represent character modifications. Asterisk (*) indicates shift in position of character in cladogram due to anatomical information from †*Coccocephalichthys*. Black bars: unambiguous changes; grey bars: ACCTRAN optimisations; white bars: DELTRAN optimisations. Blue: endocast; red: brain and cranial nerves; pink: myelencephalic sheet. Arrow indicates anterior direction for 3D renders. Insets show transverse or sagittal sections through the relevant portions of the brain, with darker orange shading indicating specific regions of interest. Images not to scale. Node ages from Giles et al.^3^.

### Patterns of brain evolution in bony fishes

The principal lineages of ray-finned fishes show substantial differences in both brain and endocavity structure (Fig. 3, Extended Data Fig. 3). Living members of early-diverging groups like cladistians and chondrosteans provide important clues about primitive brain anatomy in actinopterygians. However, both groups show extreme morphological specialisations resulting from their long independent evolutionary histories. As a stem actinopterygian separated from the common ancestor of all living species by tens—rather than hundreds—of millions of years^3,29,52,53^, †*Coccocephalichthys* provides unique information bearing on primitive brain anatomy in ray-finned fishes and sequences of change within the group. Most notably, the brain of †*Coccocephalichthys* allows us to clarify neurological synapomorphies of the ray-finned fish total group (i.e. the living radiation and all closely related fossil taxa) and crown group (i.e. the living radiation only), summarised in Fig. 3. An everted forebrain, the principle neuroanatomical feature of ray-finned fishes, is absent in †*Coccocephalichthys*, indicating that this feature originated in more crownward portions of the actinopterygian stem. Absence of this feature also nullifies the hypothesis that forebrain eversion in actinopterygians arose due to developmental constraints associated with small body size in Devonian members of the group^54,55^.

The presence of partially developed inferior lobes of the hypothalamus in †*Coccocephalichthys* challenges the current assumption that the absence of this diencephalic outgrowth in *Polypterus* represents a primitive condition for crown ray-finned fishes^1,56^.

Presence of this feature in a stem actinopterygian suggests an alternative scenario where it arose deep on the ray-finned fish stem, and was retained by actinopterans and lost in cladistians, before developing fully in neopterygians^56^. †*Coccocephalichthys* also provides evidence that the myelencephalic gland of holosteans and chondrosteans can trace its origins to a feature present in stem actinopterygians. The myelencephalic gland is a hematopoietic (blood-generating) structure enclosed within the endocranial cavity of non-teleost actinopterans, where it either overlies (lepisosteids) or embraces (*Amia*, chondrosteans) the myelencephalon^44,57^. In *Polypterus*, the meningeal tissue occupying the same region as the myelencephalic gland of other taxa is differentiated and highly vascularised, and is identified as the cisterna spinobulbaris^42,43^.

†*Coccocephalichthys* bears a similar membranous structure overlying the rhombencephalon at the level of the vagal nerves, which we consider to be homologous to the cisterna spinobulbaris of *Polypterus*. On this basis, we argue that modified rhombencephalic meningeal tissues are a general feature of ray-finned fishes, with subsequent modifications in holosteans and chondrosteans as a well-developed myelencephalic gland.

The brain of †*Coccocephalichthys* aids in discerning which neuroanatomical features of deeply-branching crown lineages are primitive versus derived, with implications for patterns of brain evolution in more nested clades (Fig. 3). These data provide remarkable corroboration that features of *Polypterus* such as the absence of intraventricular projections and the presence of a poorly differentiated corpus cerebelli represent primitive actinopterygian conditions. However, †*Coccocephalichthys* suggests that perhaps the most conspicuous external aspect of neuroanatomy in *Polypterus* might be apomorphic. Like sarcopterygians, *Polypterus* has an enlarged telencephalon^9^, in contrast to the small structure in actinopterans^38,39^ and chondrichthyans^35^. Distribution among extant taxa suggests the shared condition in *Polypterus* and sarcopterygians may be an osteichthyan feature^42^ lost in actinopterans. However, the absence of enlarged telencephalon in †*Coccocephalichthys* makes it more parsimonious to infer the convergent origin of similar geometries in *Polypterus* and sarcopterygians. At the same time, †*Coccocephalichthys* suggests that an apparent specialisation of *Polypterus* might in fact be a more general feature of ray-finned fishes. *Polypterus* is unique among extant jawed vertebrates in having an invaginated corpus cerebelli, a condition most parsimoniously interpreted as a specialisation of that lineage^9,11,58^. However, the corpus cerebelli of †*Coccocephalichthys* also seems to be formed as an invagination of the dorsal surface of the rhombencephalic region of the brain, matching the arrangement of *Polypterus*. Independent gains within both lineages, or a single gain at the base of actinopterygians followed by a loss in actinopterans, represent equally parsimonious scenarios. It is not possible to select between these alternatives in the absence of additional information on brain structure in other early actinopterygians.

### The utility of fossil brains

†*Coccocephalichthys* reinforces results from studies of neural structures in fossil arthropods^12–16^ that highlight the importance of fossil brains for patterns of neuroanatomical evolution in groups with deep evolutionary divergences. Beyond representing preservational curiosities, fossilised brains provide otherwise inaccessible trait data with implications for patterns of phylogenetic relationships and character polarity. We anticipate that preservation of neural tissue in fossil fishes is likely to be more common than widely thought, with assumptions of non-preservation leading to potentially valuable information being overlooked. A careful survey of fish material from taphonomically promising horizons has potential to yield novel anatomical information bearing on the evolution of brain structural diversity within the principal clade of aquatic vertebrates.

## Methods

### Material examined

†*Coccocephalichthys wildi* is known from a single specimen (Manchester Museum, Wild Collection, 12451) from the roof of the Mountain Fourfoot Mine, Carre Heys, Trawden, Lancashire, UK. Accounts of its anatomy are given by Watson^60^, Poplin^27^, and Poplin & Véran^61^. Other three-dimensionally preserved actinopterygians hosted in nodules from this area include *Trawdenia planti* and *Mesonichthys aitkeni*; these are all thought to derive from the so-called “Soapstone Bed.” This horizon lies within the Pennine Lower Coal Measures above the Bullion Coal (= Upper Foot Coal) and the Mountain 1.2 m Coal (= Lower Mountain Coal), but below the Ardley Seam (=Arley Coal)^26,62,63^. This is within the Langsettian regional substage, which correlates with the upper part of the Bashkirian stage of the international timescale^64^.

### Diffusible Iodine-based contrast enhancement (diceCT)

Comparative specimens of *Squalus acanthias* (University of Michigan Museum of Zoology [UMMZ] uncatalogued), *Polypterus senegalus* (UMMZ 195008), *Amia calva* (UMMZ 235291) and *Acipenser fulvicens* (UMMZ 219456) were prepared for diceCT by submerging specimens in 1.25% Lugol’s solution (25g IM_2_ + 50g KI for every 2L of water) for roughly 14 days prior to scanning. DiceCT data for a specimen of *Lepidosiren paradoxa* (UF:FISH:129826) from the Florida Museum of Natural History Ichthyology Collection was obtained from Morphosource (ark:/87602/m4/M167969).

### X-ray computed tomography

†*Coccocephalichthys wildi* and extant comparative material were scanned at the CTEES facility of the Department of Earth and Environmental Sciences, University of Michigan, using a Nikon XT H 225ST μCT scanner. The scan for †*Coccocephalichthys wildi* was set with 120 kV energy, 125 μA current and using a 0.5 mm copper filter. Eight frames were acquired for each projection, with an exposure time of 2.83 seconds, and the option for minimising ring artifacts was selected. Effective pixel size was 15.35 μm and geometric magnification = 13.031. Parameters for extant comparative material (*Squalus acanthias, Polypterus senegalus, Acipenser brevirostrum*, and *Amia calva*) can be accessed through the Supplemental Material.

## Acknowledgements

We thank David Gelsthorpe and Lindsay Loughtman (Manchester Museum) for collections access, and Ramon Nagesan and Randall Singer (UMMZ) for assistance with extant material. Lauren Simonitis and Kayla Hall (Friday Harbor Labs) are thanked for providing comparative material of *Squalus*. Alessio Capobianco, Jesús Díaz-Cruz, Carlos Mauricio Peredo provided feedback on an earlier version of this contribution, and Richard Dearden assisted with Blender. S.G. was supported by a Royal Society Dorothy Hodgkin Research Fellowship (DH160098). This study includes data produced in the CTEES facility at University of Michigan, supported by the Department of Earth and Environmental Sciences and College of Literature, Science, and the Arts.

## Author contributions

The project was conceived by M.F. and S.G. CT scanning was carried out by M.F. and R.F., with staining of extant material by R.F. and M.K. Segmentation of CT data was performed by M.F., S.G., D.G., and R.F. M.F, S.G., and R.F wrote the manuscript, with comments from all authors.

## Data availability

The fossil described in this study is deposited in the collections of the Manchester Museum and the extant specimens in the University of Michigan Museum of Zoology. The reconstructed .TIFF stack, segmented Mimics file and .PLY files for †*Coccocephalichthys wildi* are available on Zenodo (10.5281/zenodo.6560305).

## Extended Data figure legends

**Extended Data Fig. 1.**
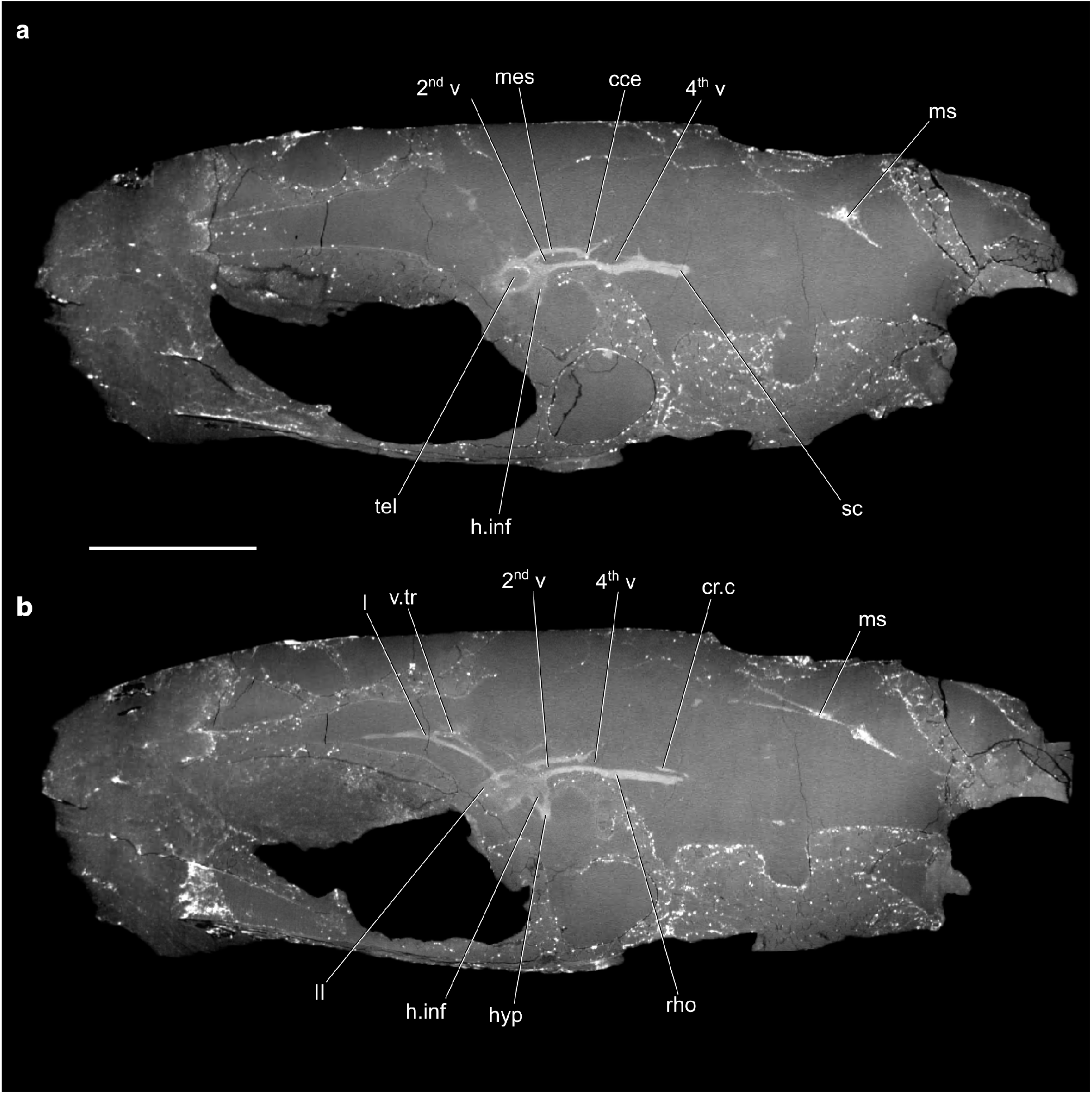
Sagittal sections through the neurocranium of †*Coccocephalichthys wildi* showing the brain and associated structures. cce, corpus cerebelli, cr.c; crista cerebellaris, h.inf, hypothalamus inferior lobes; hyp, hypophysis; mes, mesencephalon; ms, myelencephalic sheet; rho, rhombencephalon; sc, spinal cord; tel, telencephalon; v.tr, velum transversum; 2^nd^ v, second ventricle; 4^th^ v, fourth ventricle; I, olfactory nerve; II, optic nerve. Scale bar = 10 mm.

**Extended Data Fig. 2.**
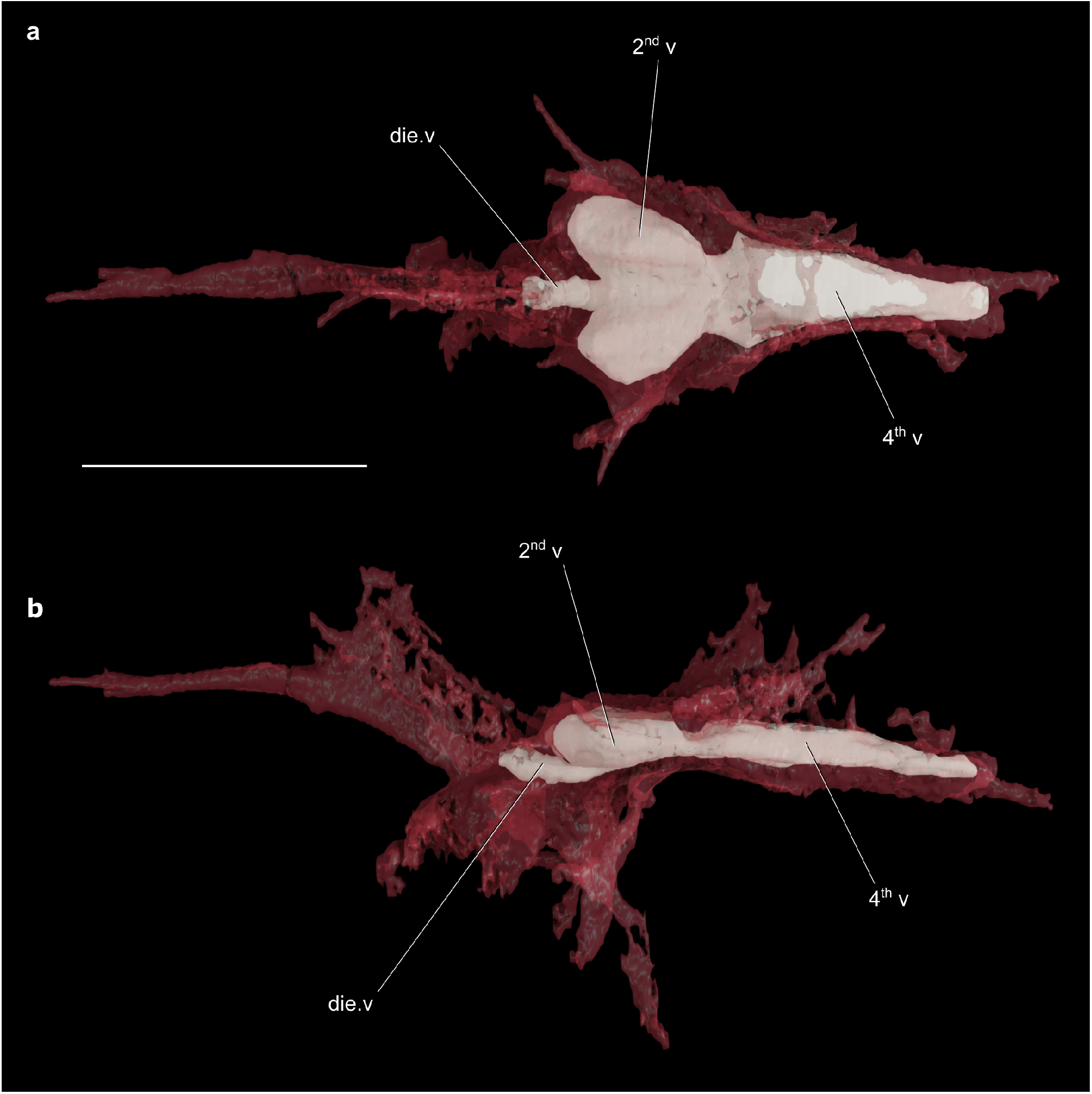
The brain of †*Coccocephalichthys wildi* (red) rendered partially transparent to show brain ventricle configuration (white). **a**, dorsal view. **b**, left lateral view. die. v, diencephalic ventricle; 2^nd^ v, second ventricle; 4^th^ v, fourth ventricle. Scale bar = 5 mm.

**Extended Data Fig. 3.**
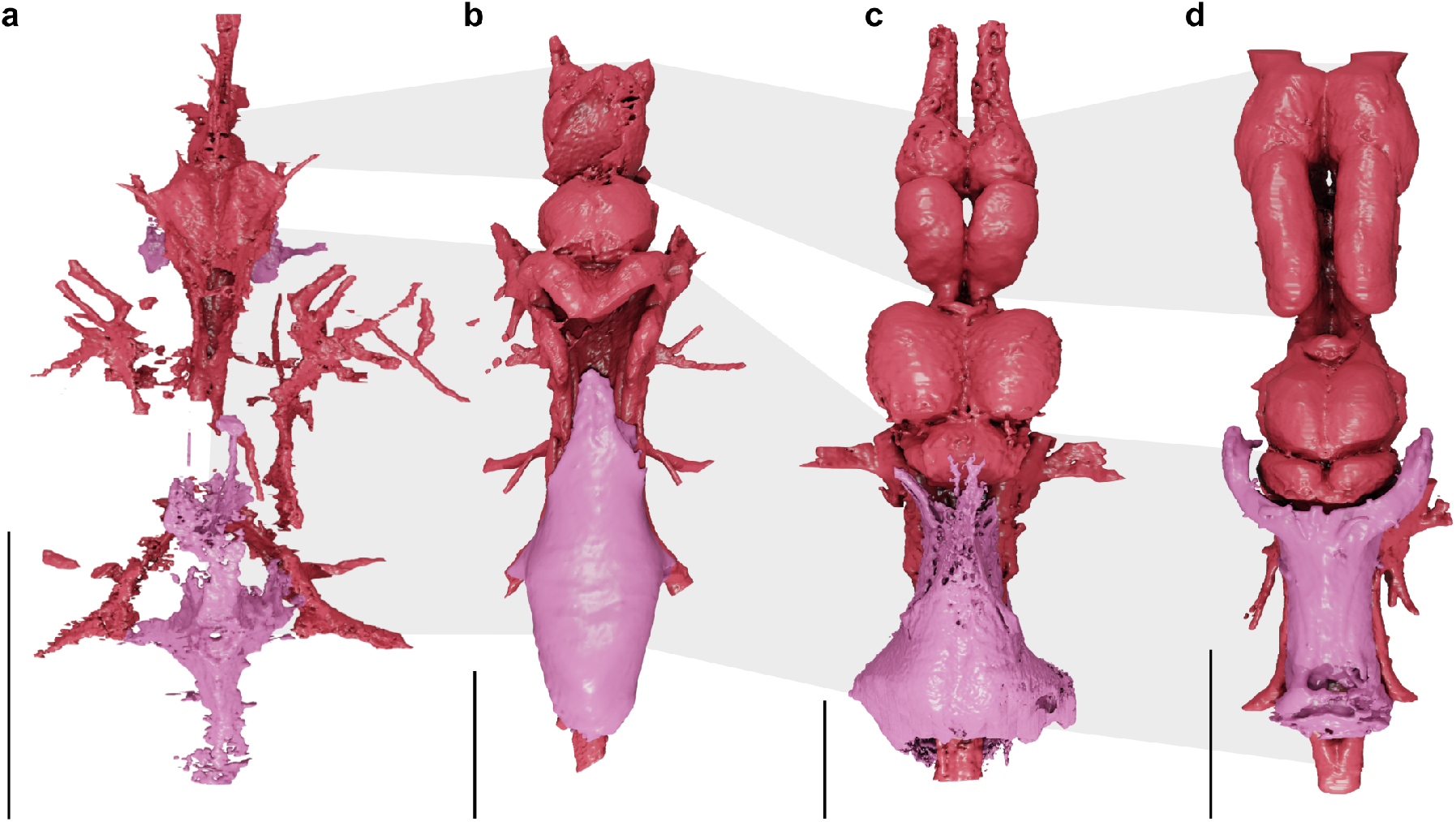
The brain (red) and myelencephalic sheet/gland (pink) of †*Coccocephalichthys wildi* and selected extant ray-finned fishes. a, **†*Coccocephalichthys wildi***. b, ***Acipenser brevirostrum***. c, ***Amia calva***. d, ***Polypterus senegalus*. Gray and white delimitations show margins between forebrain, midbrain and hindbrain across all taxa. Scale bar = 10 mm**.

**Extended Data Fig. 4.**
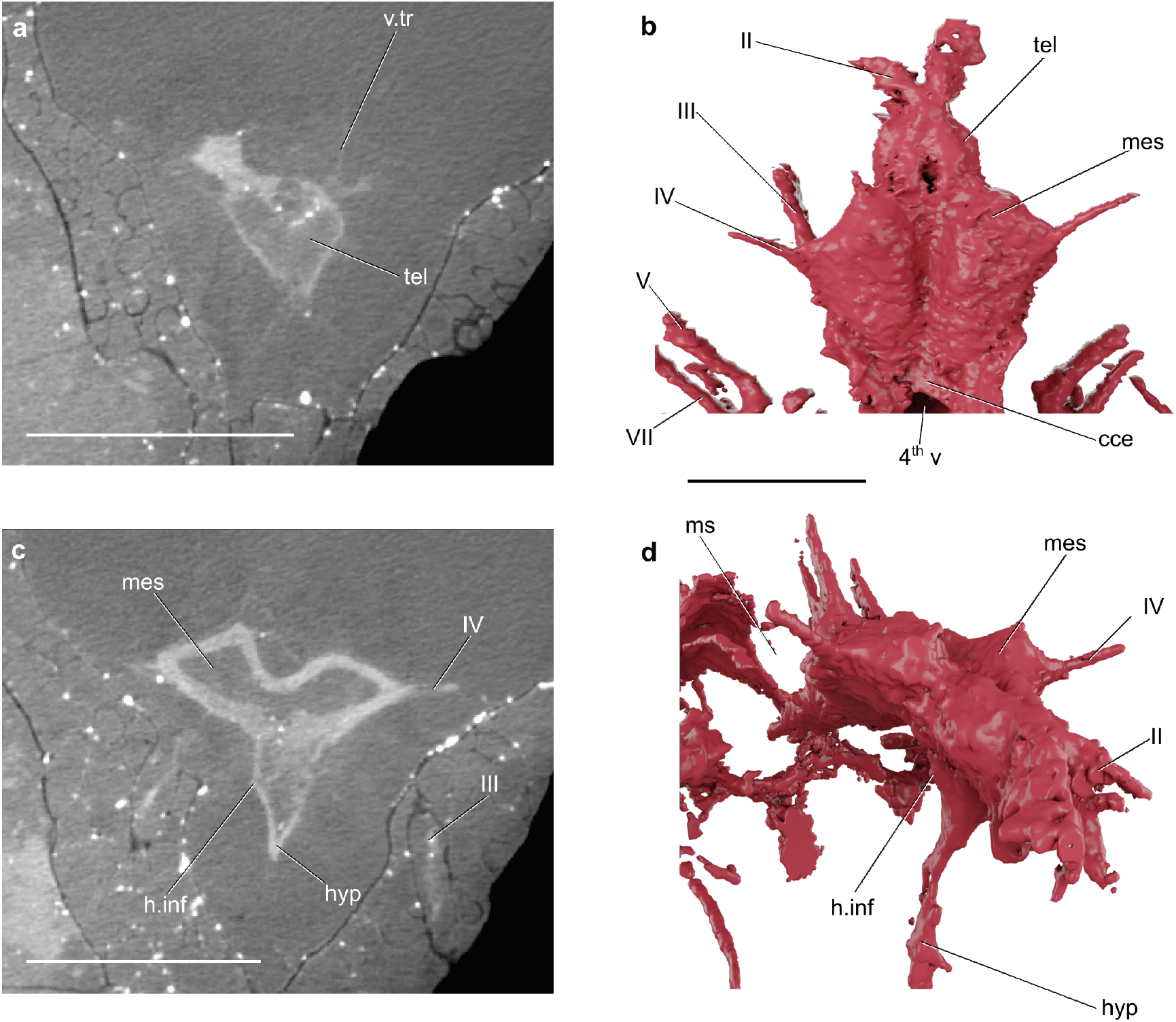
Transverse sections and renders of the brain of †*Coccocephalichthys wildi*. **a, b**, the telencephalon. **c, d**, the mesencephalon and hypophysis. cce, corpus cerebellum; h.inf, inferior lobe of the hypothalamus; hyp, hypophysis; tel, telencephalon; mes, mesencephalon; ms, mesencephalic sheet; v. tr, velum transversum; 4^th^ v, fourth ventricle; II, optic nerve; III, oculomotor nerve; IV, trochlear nerve, V, trigeminal nerve; VII, facial nerve. Dorsal portion of forebrain and velum transversum digitally removed in renders. Scale bar in a, c = 2.5 mm; scale bar in b, d = 5 mm.

**Extended Data Fig. 5.**
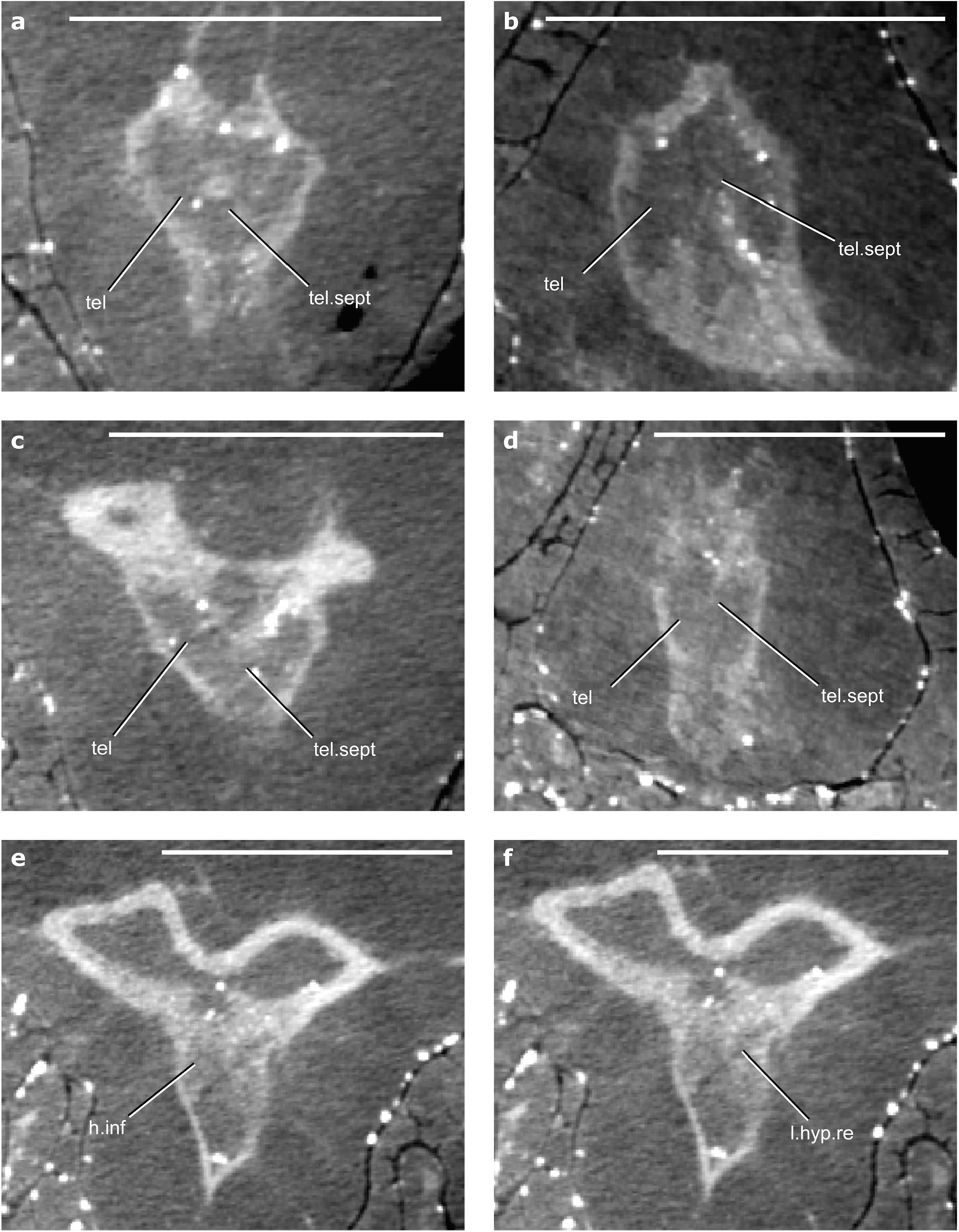
Sections through the brain of †*Coccocephalichthys wildi*. **a**, transverse section through the anterior portion of the telencephalon. **b**, axial section through the ventral portion of the telencephalon. **c**, transverse section through the posterior portion of the telencephalon. **d**, axial section through the dorsal portion of the telencephalon. **e**, transverse section through the anterior portion of the hypothalamus inferior lobes. **f**, transverse section through the posterior portion of the hypothalamus inferior lobes. h.inf, inferior lobe of the hypothalamus; l.hyp.re, lateral hypothalamic recess; tel, telencephalon; tel.sept, telencephalic septum. Scale bar = 2 mm.

**Extended Data Fig. 6.**
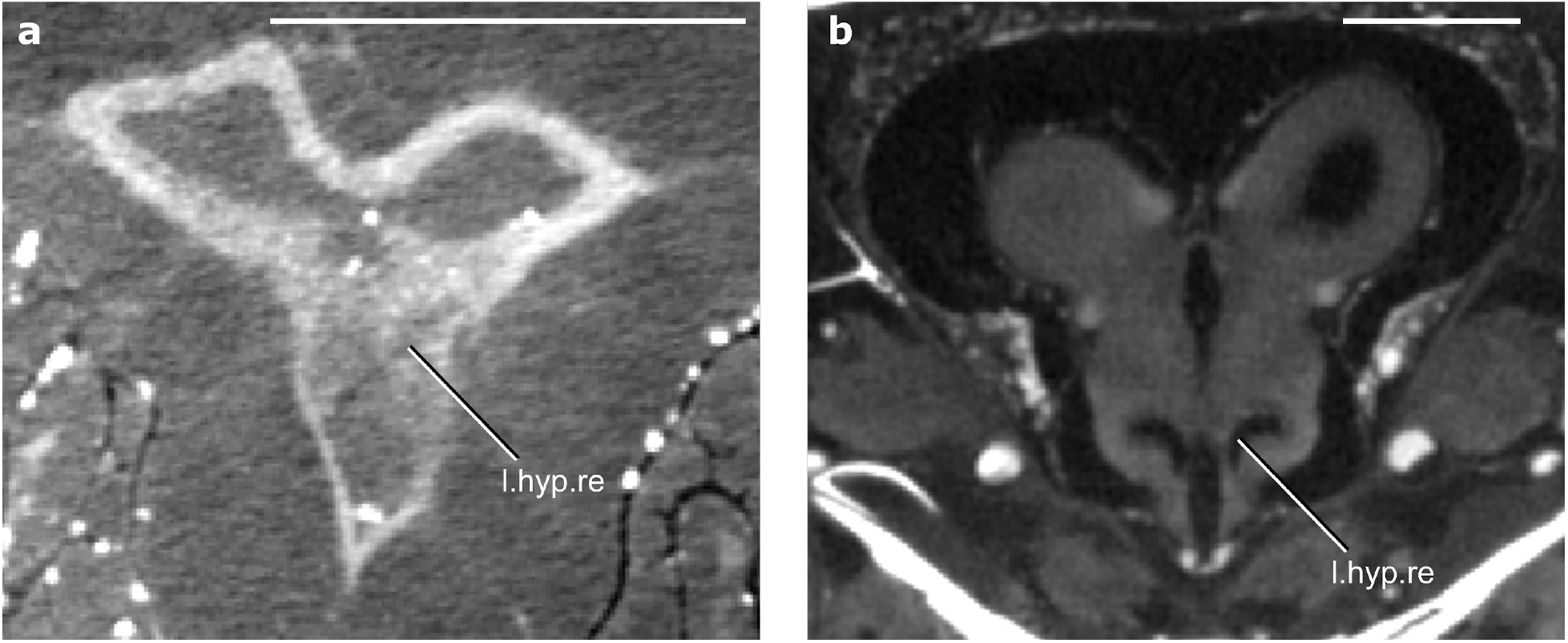
Sections through the brain of †*Coccocephalichthys wildi* and *Amia calva*. **a**, transverse section through the diencephalon and mesencephalon of *Coccocephalichthys wildi*. **b**, transverse section through the diencephalon and mesencephalon of *Amia calva*. l.hyp.re, lateral hypothalamic recess. Scale bar = 2 mm.

## References

1. Nieuwenhuys, R., ten Donkelaar, H. J. & Nicholson, C. The Meaning of It All. in The Central Nervous System of Vertebrates: Volume 1 / Volume 2 / Volume 3 (eds. Nieuwenhuys, R., ten Donkelaar, H. J. & Nicholson, C.) 2135–2195 (Springer, 1998). doi:10.1007/978-3-642-18262-4_24.

2. Friedman, M. The early evolution of ray-finned fishes. Palaeontology 58, 213–228 (2015).

3. Giles, S., Xu, G. H., Near, T. J. & Friedman, M. Early members of ‘living fossil’ lineage imply later origin of modern ray-finned fishes. Nature 549, 265–268 (2017).

4. Moodie, R. L. A new fish brain from the coal measures of Kansas, with a review of other fossil brains. J. Comp. Neurol. 25, 135–181 (1915).

5. Nielsen, E. Studies on Triassic Fishes: Glaucolepis and Boreosomus. 1. (Reitzel, 1942).

6. Giles, S. & Friedman, M. Virtual reconstruction of endocast anatomy in early ray-finned fishes (Osteichthyes, Actinopterygii). J. Paleontol. 88, 636–651 (2014).

7. Lu, J., Giles, S., Friedman, M., den Blaauwen, J. L. & Zhu, M. The oldest actinopterygian highlights the cryptic early history of the hyperdiverse ray-finned fishes. Curr. Biol. 26, 1602–1608 (2016).

8. Edinger, T. Recent Advances in Paleoneurology. Prog. Brain Res. 6, 147–160 (1964).

9. Nieuwenhuys, R. Brachiopterygian Fishes. in The Central Nervous System of Vertebrates: Volume 1 / Volume 2 / Volume 3 (eds. Nieuwenhuys, R., ten Donkelaar, H. J. & Nicholson, C.) 655–699 (Springer, 1998). doi:10.1007/978-3-642-18262-4_13.

10. Anzeiger, A. & Page, P. Beitrag zur anatomie des Zentralnervensystems und des geruchsorgans von Polypterus bichir. Aus Dem Anat. Inst. Univ. Freibg. 308–324 (1887).

11. Ikenaga, T. et al. Morphological analysis of the cerebellum and its efferent system in a basal actinopterygian fish, Polypterus senegalus. J. Comp. Neurol. 530, 1231–1246 (2022).

12. Ma, X., Hou, X., Edgecombe, G. D. & Strausfeld, N. J. Complex brain and optic lobes in an early Cambrian arthropod. (2012) doi:10.1038/nature11495.

13. Cong, P., Ma, X., Hou, X., Edgecombe, G. D. & Strausfeld, N. J. Brain structure resolves the segmental affinity of anomalocaridid appendages. (2014) doi:10.1038/nature13486.

14. Ma, X., Cong, P., Hou, X., Edgecombe, G. D. & Strausfeld, N. J. An exceptionally preserved arthropod cardiovascular system from the early Cambrian. Nat. Commun. 5, 3560 (2014).

15. Edgecombe, G. D., Ma, X. & Strausfeld, N. J. Unlocking the early fossil record of the arthropod central nervous system. Philos. Trans. R. Soc. B Biol. Sci. 370, 20150038 (2015).

16. Strausfeld, N. J., Ma, X. & Edgecombe, G. D. Fossils and the evolution of the arthropod brain. Curr. Biol. 26, R989–R1000 (2016).

17. Johanson, Z. Placoderm branchial and hypobranchial muscles and origins in jawed vertebrates. J. Vertebr. Paleontol. 23, 735–749 (2003).

18. Trinajstic, K., Marshall, C., Long, J. & Bifield, K. Exceptional preservation of nerve and muscle tissues in Late Devonian placoderm fish and their evolutionary implications. Biol. Lett. 3, 197–200 (2007).

19. Pradel, A. et al. Skull and brain of a 300-million-year-old chimaeroid fish revealed by synchrotron holotomography. Proc. Natl. Acad. Sci. U. S. A. 106, 5224–8 (2009).

20. Maldanis, L. et al. Heart fossilization is possible and informs the evolution of cardiac outflow tract in vertebrates. eLife 5, (2016).

21. Braford, M. R. Stalking the everted telencephalon: comparisons of forebrain organization in basal ray-finned fishes and teleosts. Brain. Behav. Evol. 74, 56–76 (2009).

22. Briscoe, S. D. & Ragsdale, C. W. Evolution of the chordate telencephalon. Curr. Biol. 29, R647– R662 (2019).

23. Nieuwenhuys, R. The development and general morphology of the telencephalon of actinopterygian fishes: synopsis, documentation and commentary. Brain Struct. Funct. 215, 141–157 (2011).

24. Nelson, J. S., Grande, T. C. & Wilson, M. V. H. Fishes of the World. (John Wiley & Sons, 2016).

25. Giles, S., Rogers, M. & Friedman, M. Bony labyrinth morphology in early neopterygian fishes (Actinopterygii: Neopterygii). J. Morphol. 279, 426–440 (2018).

26. Coates, M. I. Endocranial preservation of a Carboniferous actinopterygian from Lancashire, UK, and the interrelationships of primitive actinopterygians. Philos. Trans. R. Soc. B Biol. Sci. 354, 435– 462 (1999).

27. Poplin, C. M. Etude de quelques paleoniscides pennsylvaniens du Kansas. Cah. Paléontol. Ed. CNRS Paris (1974).

28. Hamel, M.-H. & Poplin, C. The braincase anatomy of Lawrenciella schaefferi, actinopterygian from the Upper Carboniferous of Kansas (USA). J. Vertebr. Paleontol. 28, 989–1006 (2008).

29. Latimer, A. E. & Giles, S. A giant dapediid from the Late Triassic of Switzerland and insights into neopterygian phylogeny. R. Soc. Open Sci. 5, 180497 (2018).

30. Argyriou, T. et al. Internal cranial anatomy of Early Triassic species of †Saurichthys (Actinopterygii: †Saurichthyiformes): implications for the phylogenetic placement of †saurichthyiforms. BMC Evol. Biol. 18, 161 (2018).

31. Choo, B., Lu, J., Giles, S., Trinajstic, K. & Long, J. A. A new actinopterygian from the Late Devonian Gogo Formation, Western Australia. Pap. Palaeontol. 5, 343–363 (2019).

32. Gignac, P. M. et al. Diffusible iodine-based contrast-enhanced computed tomography (diceCT): an emerging tool for rapid, high-resolution, 3-D imaging of metazoan soft tissues. J. Anat. 228, 889– 909 (2016).

33. Pradel, A., Maisey, J. G., Mapes, R. H. & Kruta, I. First evidence of an intercalar bone in the braincase of “palaeonisciform” actinopterygians, with a virtual reconstruction of a new braincase of Lawrenciella Poplin, 1984 from the Carboniferous of Oklahoma. Geodiversitas 38, 489–504 (2016).

34. Friedman, M. & Giles, S. Actinopterygians: the ray-finned fishes—an explosion of diversity. in Evolution of the vertebrate ear : evidence from the fossil record (eds. Clack, J. A., Fay, R. R. & Popper, A. N.) 17–49 (Springer International Publishing, 2016). doi:10.1007/978-3-319-46661-3_2.

35. Smeets, W. J. A. J. Cartilaginous Fishes. in The Central Nervous System of Vertebrates: Volume 1 / Volume 2 / Volume 3 (eds. Nieuwenhuys, R., ten Donkelaar, H. J. & Nicholson, C.) 551–654 (Springer, 1998). doi:10.1007/978-3-642-18262-4_12.

36. Nieuwenhuys, R. Lungfishes. in The Central Nervous System of Vertebrates: Volume 1 / Volume 2 / Volume 3 (eds. Nieuwenhuys, R., ten Donkelaar, H. J. & Nicholson, C.) 939–1006 (Springer, 1998). doi:10.1007/978-3-642-18262-4_16.

37. Nieuwenhuys, R. The Coelacanth Latimeria chalumnae. in The Central Nervous System of Vertebrates: Volume 1 / Volume 2 / Volume 3 (eds. Nieuwenhuys, R., ten Donkelaar, H. J. & Nicholson, C.) 1007–1043 (Springer, 1998). doi:10.1007/978-3-642-18262-4_17.

38. Nieuwenhuys, R. Chondrostean Fishes. in The Central Nervous System of Vertebrates: Volume 1 / Volume 2 / Volume 3 (eds. Nieuwenhuys, R., ten Donkelaar, H. J. & Nicholson, C.) 701–757 (Springer, 1998). doi:10.1007/978-3-642-18262-4_14.

39. Meek, J. & Nieuwenhuys, R. Holosteans and Teleosts. in The Central Nervous System of Vertebrates: Volume 1 / Volume 2 / Volume 3 (eds. Nieuwenhuys, R., ten Donkelaar, H. J. & Nicholson, C.) 759–937 (Springer, 1998). doi:10.1007/978-3-642-18262-4_15.

40. Northcutt, R. G. Forebrain evolution in bony fishes. Brain Res. Bull. 75, 191–205 (2008).

41. Morona, R., López, J. M., Northcutt, R. G. & González, A. Comparative analysis of the organization of the cholinergic system in the brains of two Holostean fishes, the Florida gar Lepisosteus platyrhincus and the bowfin Amia calva. Brain. Behav. Evol. 81, 109–142 (2013).

42. Jarvik, E. Basic structure and evolution of vertebrates. (Academic Press, 1980).

43. Bjerring, H. C. Facts and thoughts on piscine phylogeny. in Evolutionary Biology of Primitive Fishes (eds. Foreman, R. E., Gorbman, A., Dodd, J. M. & Olsson, R.) 31–57 (Springer US, 1985). doi:10.1007/978-1-4615-9453-6_3.

44. Chandler, A. C. On a lymphoid structure lying over the myelencephalon of Lepisosteus. Univ. Calif. Publ. Zool. 9, 85–104 (1911).

45. Coates, M. I. Actinopterygians from the Namurian of Bearsden, Scotland, with comments on early actinopterygian neurocrania. Zool. J. Linn. Soc. 122, 27–59 (1998).

46. Fine, M. L., Horn, M. H. H. & Cox, B. Acanthonus armatus, a deep-sea teleost fish with a minute brain and large ears. Proc. R. Soc. Lond. B Biol. Sci. 230, 257–265 (1987).

47. Herzog, H., Klein, B. & Ziegler, A. Form and function of the teleost lateral line revealed using three-dimensional imaging and computational fluid dynamics. J. R. Soc. Interface 14, 20160898 (2017).

48. Rowe, T. B., Macrini, T. E. & Luo, Z.-X. Fossil evidence on origin of the mammalian brain. Science 332, 955–957 (2011).

49. Neubauer, S. Endocasts: possibilities and limitations for the interpretation of human brain evolution. Brain. Behav. Evol. 84, 117–134 (2014).

50. Clement, A. M., Nysjö, J., Strand, R. & Ahlberg, P. E. Brain -Endocast relationship in the Australian lungfish, Neoceratodus forsteri, elucidated from tomographic data (Sarcopterygii: Dipnoi). PLoS ONE 10, (2015).

51. Watanabe, A. et al. Are endocasts good proxies for brain size and shape in archosaurs throughout ontogeny? J. Anat. 234, 291–305 (2019).

52. Figueroa, R. T., Friedman, M. & Gallo, V. Cranial anatomy of the predatory actinopterygian Brazilichthys macrognathus from the Permian (Cisuralian) Pedra de Fogo Formation, Parnaíba Basin, Brazil. J. Vertebr. Paleontol. 39, e1639722 (2019).

53. Stack, J. & Gottfried, M. D. A new, exceptionally well-preserved Permian actinopterygian fish from the Minnekahta Limestone of South Dakota, USA. J. Syst. Palaeontol. 19, 1271–1302 (2021).

54. Striedter, G. F. & Northcutt, R. G. Head size constrains forebrain development and evolution in ray-finned fishes. Evol. Dev. 8, 215–222 (2006).

55. Folgueira, M. et al. Morphogenesis underlying the development of the everted teleost telencephalon. Neural Develop. 7, 212 (2012).

56. Schmidt, M. Evolution of the hypothalamus and inferior lobe in ray-finned fishes. Brain. Behav. Evol. 95, 302–316 (2020).

57. van der Horst, C. J. The myelencephalic gland of Polyodon, Acipenser and Amia. K. Akad. Van Wet. Te Amst. Proc. Sect. Sci. 28, 432–442 (1925).

58. Graña, P., Folgueira, M., Huesa, G., Anadón, R. & Yáñez, J. Immunohistochemical distribution of calretinin and calbindin (D-28k) in the brain of the cladistian Polypterus senegalus. J. Comp. Neurol. 521, 2454–2485 (2013).

## Methods references

59. Lankester, E. R. & Ridewood, W. G. Guide to the gallery of fishes. (British Museum (Natural History). Department of Zoology, 1908).

60. Watson, D. M. S. The structure of certain palaeoniscids and the relationships of that group with other bony fish. Proc. Zool. Soc. Lond. 95, 815–870 (1925).

61. Poplin, C. M. & Véran, M. A revision of the actinopterygian fish Coccocephalus wildi from the Upper Carboniferous of Lancashire. Spec. Pap. Palaeontol. 52, 7–29 (1996).

62. Coates, M. I. & Tietjen, K. ‘This strange little palaeoniscid’: a new early actinopterygian genus, and commentary on pectoral fin conditions and function. Earth Environ. Sci. Trans. R. Soc. Edinb. 109, 15–31 (2018).

63. Hough, E. Geology of the Burnley area (SD82NW and SD83SW). (2004).

64. Waters, C. N. et al. A revised correlation of carboniferous rocks in the British Isles. (Geological Society of London, 2011). doi:10.1144/SR26.

