## Supplementary Information for "Exceptional fossil preservation and evolution of the ray-finned fish brain"

**Phylogenetic placement of †*Coccocephalichthys wildi***

†*Coccocephalichthys wildi* has been included in a number of phylogenetic analyses investigating the relationships of early ray-finned fishes[1–11]. It was originally placed in the actinopterygian crown[1], but all subsequent analyses resolve it on the actinopterygian stem, sometimes within a large polytomy including most other late Palaeozoic ray-finned fishes. A detailed redescription of †*Coccocephalichthys* based on new μCT data, in conjunction with an updated phylogenetic analysis, is currently in preparation (S.G., R.F., M.F.), although we do not anticipate this leading to a change in the phylogenetic placement of the taxon.

The cranial morphology of †*Coccocephalichthys wildi* is comparable to other late Palaeozoic taxa that are well-within the ray-finned fish stem. The clearest comparisons are with the Carboniferous †*Kentuckia*[12], †*Kansasiella*[13], and †*Lawrenciella*[13–15], all of which preserve information on the endocranium. In common with these taxa, †*Coccocephalichthys* has a completely enclosed spiracular canal, a posteroventrally-directed hyoid facet, and open vestibular fontanelles continuous with the oticoocippital fissure. As in †*Kentuckia* and †*Lawrenciella*, the aortic canal of †*Coccocephalichthys* is pierced by a single midline opening, and as in †*Kentuckia* the divergence of the lateral dorsal aortae is not enclosed within the aortic canal and the parasphenoid is flat rather than dorsally inflected below the orbit. †*Coccocephalichthys* also shares other features in common with a subset of the above taxa, as well as with some Devonian and stratigraphically younger taxa.

†*Coccocephalichthys* is excluded from the crown by several successive nodes on the basis of the following ambiguous character optimisations (taken from ref. 5):

- Character 42: bone carrying otic portion of lateral line canal extends past posterior margin of parietals
- Character 89: two infradentaries
- Character 94: coronoid process formed by suragular
- Character 100: comineralised palatoquadrate ossifications
- Character 156: braincase comprises single ossification
- Character 171: parasphenoid terminates at/anterior to ventral otic fissure
- Character 178: aortic notch absent from parasphenoid
- Character 43: two pairs of extrascapulars
- Character 69: maxilla contributes to posterior margin of cheek
- Character 153: dorsal aorta not pierced by canal/s for exit of eff.a.1
- Character 175: parasphenoid teeth small

**CT scanning parameters for all specimens**

| | Energy (kV) | Current (μA) | Exposure (S) | Filter | Filter (mm) | Voxel (mm) | Frames/ Projection | Minimize Ring Artifact |
|---|---|---|---|---|---|---|---|---|
| ***Coccocephalichthys wildi*** (MM12451) | 120 | 125 | 2.83 | Cu | 0.5 | 0.01535 | 8 | YES |
| ***Squalus acanthias*** (UMMZ uncatalog.) | 160 | 120 | 1.42 | NA | - | 0.06243 | 2 | YES |
| ***Polypterus senegalus*** (UMMZ 195008) | 100 | 172 | 0.25 | NA | - | 0.01733 | 16 | YES |
| ***Acipenser brevirostrum*** (UMMZ 219456) | 85 | 200 | 0.25 | NA | - | 0.03236 | 16 | YES |
| ***Amia calva*** (UMMZ 235291) | 85 | 200 | 0.25 | NA | - | 0.03451 | 16 | YES |

1. Coates, M. I. Endocranial preservation of a Carboniferous actinopterygian from Lancashire, UK, and the interrelationships of primitive actinopterygians. *Philos. Trans. R. Soc. B Biol. Sci.* 354, 435–462 (1999).
2. Cloutier, R. & Arratia, G. Early diversification of actinopterygians. in *The origin and early radiation of vertebrates: honoring Hans-Peter Schultze* (Verlag Dr. Friedrich Pfeil, 2004).
3. Giles, S., Darras, L., Clément, G., Blieck, A. & Friedman, M. An exceptionally preserved Late Devonian actinopterygian provides a new model for primitive cranial anatomy in ray-finned fishes. *Proc. R. Soc. B Biol. Sci.* **282**, (2015).
4. Giles, S., Xu, G. H., Near, T. J. & Friedman, M. Early members of 'living fossil' lineage imply later origin of modern ray-finned fishes. *Nature* **549**, 265–268 (2017).
5. Latimer, A. E. & Giles, S. A giant dapediid from the Late Triassic of Switzerland and insights into neopterygian phylogeny. *R. Soc. Open Sci.* **5**, 180497 (2018).
6. Wilson, C. D., Pardo, J. D. & Anderson, J. S. A primitive actinopterygian braincase from the tournaisian of Nova Scotia. *R. Soc. Open Sci.* **5**, (2018).
7. Argyriou, T. *et al.* Internal cranial anatomy of Early Triassic species of †*Saurichthys* (Actinopterygii: †Saurichthyiformes): implications for the phylogenetic placement of †saurichthyiforms. *BMC Evol. Biol.* **18**, 161 (2018).
8. Choo, B., Lu, J., Giles, S., Trinajstic, K. & Long, J. A. A new actinopterygian from the Late Devonian Gogo Formation, Western Australia. *Pap. Palaeontol.* **5**, 343–363 (2019).
9. Figueroa, R. T., Friedman, M. & Gallo, V. Cranial anatomy of the predatory actinopterygian *Brazilichthys macrognathus* from the Permian (Cisuralian) Pedra de Fogo Formation, Parnaíba Basin, Brazil. *J. Vertebr. Paleontol.* e1639722 (2019) doi:10.1101/540310.
10. Figueroa, R. T., Weinschütz, L. C. & Friedman, M. The oldest Devonian circumpolar ray-finned fish? *Biol. Lett.* **17**, 20200766 (2021).
11. Stack, J. & Gottfried, M. D. A new, exceptionally well-preserved Permian actinopterygian fish from the Minnekahta Limestone of South Dakota, USA. *J. Syst. Palaeontol.* **19**, 1271–1302 (2021).

12. Rayner, D. H. On the cranial structure of an early palaeoniscid, *Kentuckia*, gen. nov. *Trans. R. Soc. Edinb.* **62**, 53–83 (1952).

13. Poplin, C. M. Etude de quelques paleoniscides pennsylvaniens du Kansas. *Cah. Paléontol. Ed. CNRS Paris* (1974).

14. Hamel, M.-H. & Poplin, C. The braincase anatomy of *Lawrenciella schaefferi* , actinopterygian from the Upper Carboniferous of Kansas (USA). *J. Vertebr. Paleontol.* **28**, 989–1006 (2008).

15. Pradel, A., Maisey, J. G., Mapes, R. H. & Kruta, I. First evidence of an intercalar bone in the braincase of "palaeonisciform" actinopterygians, with a virtual reconstruction of a new braincase of *Lawrenciella* Poplin, 1984 from the Carboniferous of Oklahoma. *Geodiversitas* **38**, 489–504 (2016).
